# Modelling double strand break susceptibility to interrogate structural variation in cancer

**DOI:** 10.1101/441832

**Authors:** Tracy J. Ballinger, Britta Bouwman, Reza Mirzazadeh, Silvano Garnerone, Nicola Crosetto, Colin A. Semple

**Affiliations:** MRC Human Genetics Unit, MRC Institute of Genetics and Molecular Medicine, University of Edinburgh, Crewe Road, Edinburgh, EH4 2XU, UK; Science for Life Laboratory, Department of Medical Biochemistry and Biophysics, Karolinska Institutet, Stockholm, Sweden

**Author notes:** These authors contributed equally to this work (Nicola Crosetto, Colin A. Semple). **Corresponding Author:** Dr Tracy Ballinger, MRC Human Genetics Unit, Institute of Genetics and Molecular Medicine, University of Edinburgh, Crewe Road, Edinburgh, EH4 2XU, UK. **Author Emails:** Tracy Ballinger Britta Bouwman Reza Mirzazadeh Silvano Garnerone Nicola Crosetto Colin Semple.

**Keywords:** Double strand break, cancer, structural variaton, chromatin, modelling

## Abstract

**Background:** Structural variants (SVs) are known to play important roles in a variety of cancers, but their origins and functional consequences are still poorly understood. Many SVs are thought to emerge via errors in the repair processes following DNA double strand breaks (DSBs) and previous studies have experimentally measured DSB frequencies across the genome in cell lines.

**Results:** Using these data we derive the first quantitative genome-wide models of DSB susceptibility, based upon underlying chromatin and sequence features. These models are accurate and provide novel insights into the mutational mechanisms generating DSBs. Models trained in one cell type can be successfully applied to others, but a substantial proportion of DSBs appear to reflect cell type specific processes. Using model predictions as a proxy for susceptibility to DSBs in tumours, many SV enriched regions appear to be poorly explained by selectively neutral mutational bias alone. A substantial number of these regions show unexpectedly high SV breakpoint frequencies given their predicted susceptibility to mutation, and are therefore credible targets of positive selection in tumours. These putatively positively selected SV hotspots are enriched for genes previously shown to be oncogenic. In contrast, several hundred regions across the genome show unexpectedly low levels of SVs, given their relatively high susceptibility to mutation. These novel ‘coldspot’ regions appear to be subject to purifying selection in tumours and are enriched for active promoters and enhancers.

**Conclusions:** We conclude that models of DSB susceptibility offer a rigorous approach to the inference of SVs putatively subject to selection in tumours.

## Background

Structural variation (SV) in tumour genomes is known to play important roles in disease progression and may be critical in driving the development of certain cancer types (1–3). However, challenges remain not only in ascertaining accurate SV calls, as evidenced by the compendium of SV calling algorithms used in many projects (4–6), but also in predicting their functional impact. Some SVs have apparently direct consequences; for example, amplification of oncogenes leading to overexpression, deletion of tumor suppressors leading to dysfunction, and translocations generating oncogenic fusion proteins (4). Reportedly indirect consequences of SVs include changes in enhancer targeting, affecting the expression of nearby genes, or “enhancer hijacking” (7). However, it remains challenging to distinguish the influences of evolutionary selection versus primary mutation rate in generating the SVs concerned.

A recent study of whole genome sequencing (WGS) data from breast tumours identified SV hotspots and putative driver SVs, but could not discern the relative contributions of mutational bias and selection underlying these hotspots (8). Resolving the influences of mutational bias versus selective forces has become critical given that both single nucleotide variant (SNV) and SV mutation rates vary widely across the genome, in parallel with replication timing and chromatin structure (9,10). In analyses of tumour SNVs, variants are routinely prioritized based on algorithms including corrections for estimates of SNV mutation rate variation (11), but analogous methods are not yet applied to SVs.

Variable rates of SVs observed across the genome are likely to be affected by differences in the efficiency of repair of DNA double strand breaks (DSBs). DSBs can be repaired by homologous recombination (HR) at the G2 and S stages of the cell cycle and, more commonly, by canonical non-homologous end joining (c-NHEJ) which operates throughout the cell cycle (12). The c-NHEJ process is error prone and has been shown to create structural variants initiating carcinogenesis (13). A third repair process, alternative NHEJ (alt-NHEJ) uses microhomology to mediate repairs when the c-NHEJ pathway is unavailable, and repair by alt-NHEJ appears to increase the rate of deletions, insertions, and translocations further (14). The efficiency of these repair processes is often dependent upon the chromatin features and nuclear organization present where the damage occurs. For example, the histone modification H3K36me3, associated with active transcription, recruits the HR pathway, while H4K20me1, a mark of highly transcribed genes, recruits components of the NHEJ pathway (15). The associations between DSB repair and the underlying chromatin landscape may therefore explain the observed correlations between tumour SV rates and chromatin structure (9).

Previous studies have also shown DSB formation to be influenced by underlying chromatin structures and genomic sequences. It has long been known that certain cytogenetically mapped loci, termed “fragile sites” undergo recurrent DSBs in cells under replicative stress and in cancer (16). More recent high throughput sequencing (HTS) based approaches have been developed to profile DSB rates more precisely within *in vitro* populations of cells (17–25). Three of these methods, BLESS (18), DSBCapture (22), and BLISS (25) are closely related and have been used to generate high-resolution maps of endogenous DSBs occurring in human cell lines, resulting in continuous data reflecting the propensities for DSBs across all chromosomes. These studies have suggested that DSBs may preferentially occur within nucleosome-depleted regions, are correlated with active promoter and enhancer histone modifications, and may associate with G-quadruplex sites (22,26). Certain studies have also suggested DSBs to be depleted in some transposon classes and enriched in some simple repeat classes, and to be unusually frequent in long, late-replicating genes (18,24). Overall, previous studies have found correlations and enrichments between DSBs and various inter-correlated chromatin and genomic features, making it difficult to accurately assess the contribution of any particular feature to DSB susceptibility. Understanding such contributions can be valuable for understanding the underlying mutational and repair mechanisms. In addition, a fuller understanding of the relative contributions of many features to DSB formation can allow reliable predictions of the expected DSB frequency in a given genomic region.

Random forests have been used to model a variety of biological phenomena because they perform well in the presence of inter-correlated input variables showing non-linear relationships. For example, they have been used to predict nuclear compartments (27), cancer SNV mutational landscapes (28), and enhancer-promoter interactions (29). In this study we construct random forest regression models to generate quantitative measures of the relative importance of a variety of matched chromatin and other features to DSB susceptibility. We use multiple, high-resolution DSB profiling datasets to compare modeling accuracy across several platforms and cell types. The cell types selected have also been extensively profiled for a variety of chromatin features by the ENCODE Project (30) and others, allowing well-matched models to be constructed for all datasets. We demonstrate that these models provide accurate estimates for the expected rate of DSBs in a given region and can be cross applied between DSB datasets. In addition the models can be used to explore tumour SV breakpoint data, to nominate novel regions putatively subject to selection in cancer.

## Results

We uniformly processed four DSB datasets from three related platforms (DSBCapture and BLISS are both based upon modifications to the BLESS protocol) and covering three different cell types, collating matched chromatin data for each. These datasets include two novel DSB mapping datasets derived from the K562 erythroleukemia and MCF7 breast cancer cell lines using the recently developed BLISS method (25) (see Methods) and two previously published DSB mapping datasets derived from the NHEK keratinocyte cell line using BLESS and DSBCapture (22) protocols. DSB frequency is defined in each dataset as the number of unique reads mapping to a given 50kb region, since each read in a DSBCapture, BLESS, or BLISS experiment represents an exposed DNA DSB end. Replicate experiments within each dataset were strongly and significantly correlated (Pearson′s r = 0.905 to 0.992, p<2.2e-16) and were combined to reduce noise, although random forest models generated from any single one of the replicates yielded very similar results (see Methods). Comparisons among DSB profiling datasets showed moderate correlations in genome-wide DSB frequency between the three cell types as expected (r = 0.351 to 0.635, p<2.2e-16), shown in Supp Figure 1. All three cell types correspond to well-characterized ENCODE cell lines, providing numerous matched chromatin and genomic features exhibiting a range of correlations to DSB (Figure 1), and are also inter-correlated themselves (Supp Figure 2).

**Figure 1:**
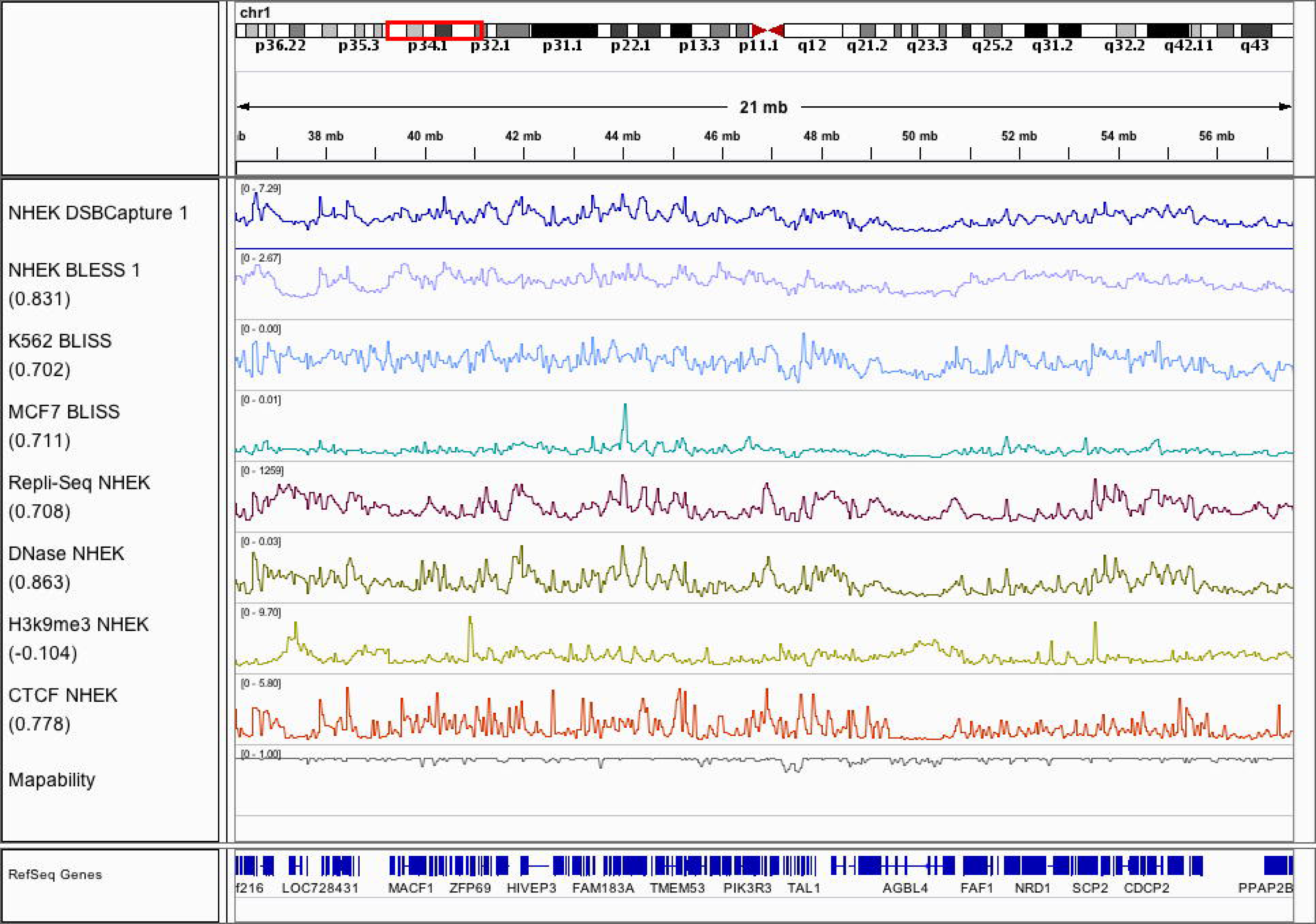
DSB frequency and genomic features display similar patterns. The tracks show DSBCapture profiles in NHEK cells, BLESS profiles in NHEK cells, BLISS in K562 cells, and BLISS in MCF7 cells. All tracks are at 50kb resolution over a representative region of chromosome 1, with a variety of chromatin and sequence features to illustrate the similarities between them. Numbers in parenthesis are the spearman′s rho between the associated track and the NHEK DSBCapture 1 dataset.

## Accurate models of genome-wide DSB frequency across cell types

We modeled DSB frequency at 50kb resolution, using the same ten matched genomic features from each cell type to construct random forest models (see Methods): open chromatin assayed by DNase-seq, POL2B binding, CTCF binding and five histone modifications assayed by ChIP-seq, replication timing assayed by Repli-seq, and RNA-seq. We also included G-quadruplex forming regions as an additional feature, since these DNA secondary structures are associated with genomic instability (31). We found strong and significant correlations between predicted and observed DSB frequency for all four datasets, with Pearson′s coefficients ranging from 0.83 to 0.92 (Figure 2). We also generated a model for the NHEK DSBCapture dataset using an extended set of 21 features, including additional histone modifications, histone variants, and nuclear compartmentalization from Hi-C data (32). This extended model resulted in better predictive results for a small fraction of the genome (Supp Figure 4, Box B), and a modestly increased genome-wide Pearson’s coefficient between predicted and observed values (11 feature model r = 0.918; 21 feature model r = 0.922). We conclude that models constructed using the 11 selected genomic features (Figure 2) provide high predictive accuracy across cell types, with additional features likely to provide only marginal gains.

**Figure 2:**
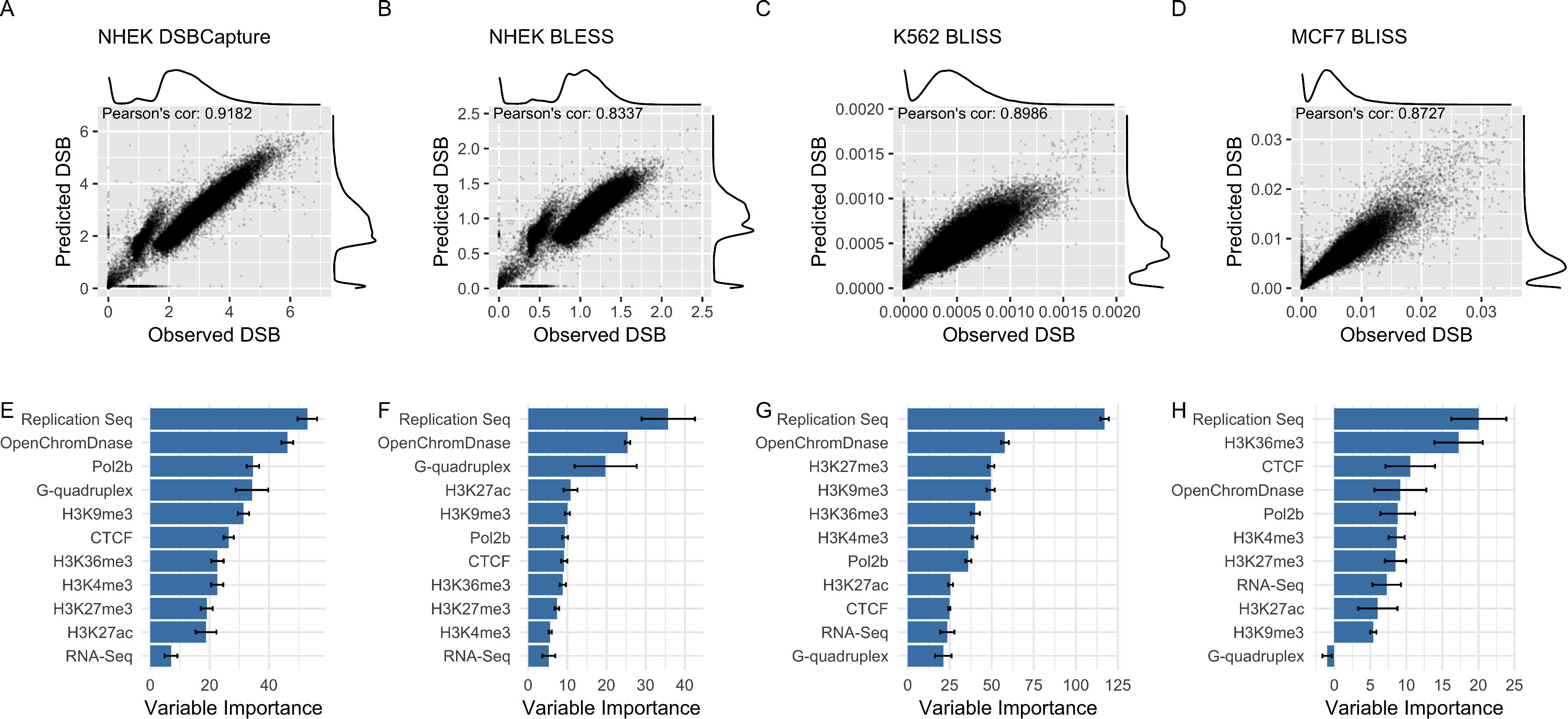
Accurate models of DSB frequency built from chromatin and sequence features. Panels A-D show random forest regression model predictions built upon eleven genomic features at 50kb resolution compared to observed DSB frequencies for four datasets: NHEK DSBCapture, NHEK BLESS, K562 BLISS, and MCF7 BLISS. The y-values reflect the sequencing depth of each dataset. The models’ predictions are all highly correlated with the observed data, as shown by the noted Pearson’s correlations (p<2.2e-16 for each dataset). Panels E-H show the predictive features ranked by variable importance, a measure of how useful a particular feature is for the model (see methods).

Variable importance metrics for these models reveal consistent trends in the most influential features in DSB frequency prediction (Figure 2,E-H). Replication timing is the most important feature across all three models with early replication associated with high DSB regions and late replication with low DSB (Figure 3C), in agreement with previous studies (33). In addition, the histone modifications H3K36me3 and H3K9me3 (demarcating active genes and gene-poor heterochromatin respectively) emerge as informative features, with H3K36m3 enriched in high DSB regions and H3K9me3 in low DSB regions (Figure 3C). This is consistent with observations that structural variants disproportionately accumulate within the early replicating, relatively gene rich regions of the genome in cancer, and are relatively depleted in late replicating heterochromatin (9,10). DNase-seq open chromatin ranks second in three datasets and fourth in the MCF7 model and is also the most important feature for predicting DSB peaks in the study of Mourad et al. (34) in which they do not include replication timing. The influence of G-quadruplex forming regions is notably variable, ranking as a relatively important feature in the NHEK datasets, but having little and no predictive value in the K562 and MCF7 datasets. RNA-seq is not a strong predictor of DSB susceptibility although DNase-seq peaks are often found at the promoter regions of active genes. This suggests that open chromatin at transcriptionally active genes and associated regulatory elements (reflected in DNase-seq, H3K4me3 and POL2B binding), rather than transcription per se, is the dominant influence on DSB frequency. CTCF binding also appears to be an informative variable, genome-wide in all models, though it binds at sites constituting a very small fraction of the genome. Given the critical roles of CTCF in chromatin architecture and regulation (32), there has been intense interest in the causes and effects of structural variants disrupting CTCF binding sites (35,36).

**Figure 3:**
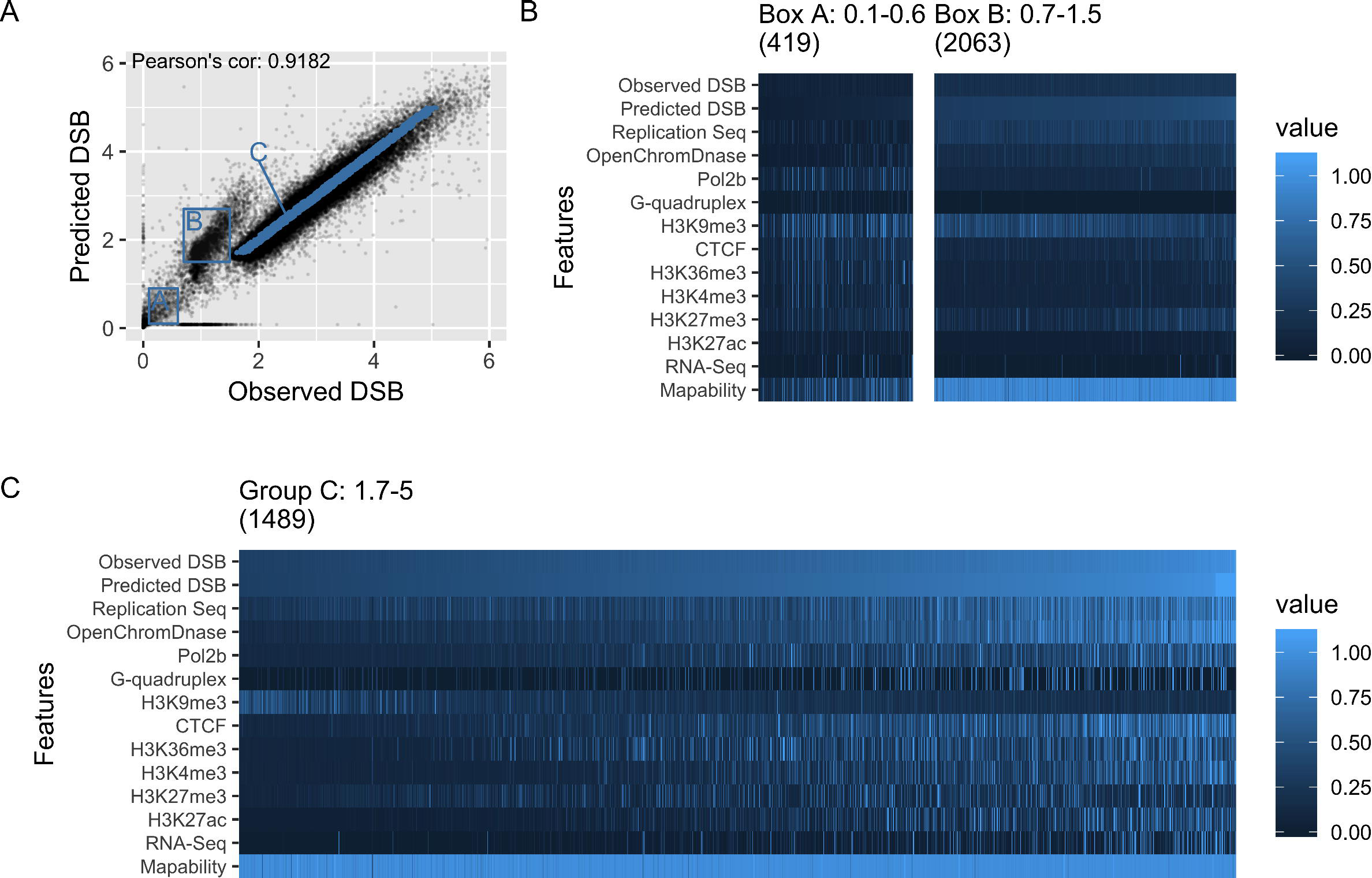
Modelling accuracy and the polarity of genomic features. A) NHEK DSBCapture 50kb regions data is split into three distinct groups with differing modelling accuracies. Panels B and C show the values of the model features for the two boxes, A and B, and for group C, which contains randomly chosen points along the spectrum of DSB frequency values for the majority of the genome. The columns are ordered by observed DSB frequency, shown on the top row, and the rows for features used to build the model (the third to second to last row) are ordered by average variable importance. The number of 50kb regions in each group is shown in parenthesis above each heatmap. Each feature was normalized, setting the 1^st^ to 99^th^ quantiles to values between 0 and 1, with high outliers (in the top percentile) set to 1.1. B) Group A has high H3K9me3 and low mappability scores, indicative of heterochromatin and repetitive sequence, while B has feature patterns that closely match low DSB values in group C. C) For most of the genome, high H3K9me3 corresponds to low DSB regions, and high, or early, replication timing values and open chromatin values signify high DSB regions.

## Influential features underlying DSB frequency differ between genomic loci and cell types

Beyond the general, genome-wide trends described above, we see differences in the behavior of certain classes of loci. These are evident as regions departing from the linear relationship between observed and predicted DSB frequency seen for the majority of the genome (Figure 3A; Supp Fig 4). Deeper exploration of the relationships between underlying genomic features and DSB frequency reveals diagnostic features for these discrepant classes. One class of loci (Figure 3, Box A) shows unusually low values for both predicted and observed DSB frequencies, and is enriched for H3K9me3 marked heterochromatin and low sequence mappability (Figure 3B). These regions are likely to correspond to repeat-rich regions near centromeres and on the short arms of acrocentric chromosomes, which are problematic for read mapping algorithms (37). Another class of H3K9me3 heterochromatin enriched loci shows higher DSB predictions than observed, in spite of high mappability values (Figure 3, Box B). This class of regions is absent in DSB datasets generated by the BLISS protocol (Figure 2), so these aberrant predictions may reflect technical and methodological differences between datasets. In any case, it is clear that model predictions may reasonably be expected to be less accurate in heterochromatic regions.

The similarities in relative variable importance across datasets (Figure 2) suggest that many features have a similar influence on DSB frequency in each of the three cell types. Thus, a model trained in one cell type might generalize well to another cell type and allow us to generate predictive DSB frequency profiles for model cell lines currently lacking high resolution DSB data. We cross-applied models and found models trained in one cell type often performed well in another (Figure 4). For example, a model trained in NHEK cells could be used to predict DSB frequencies in K562 cells (inputting K562 genomic features) with high accuracy (Pearson’s r = 0.85 correlation; Figure 4). This offers a substantial improvement over the base correlation (r = 0.63) between NHEK and K562 observed DSB profiles. We measured the correlation of observed and predicted DSB frequencies across all nine model and feature combinations and always found correlations (r = 0.58 to 0.85) that improved on the base correlations (r = 0.38 to 0.63) seen between the observed DSB datasets (Figure 4). These improvements echo the similarities in variable importance between cell types (Figure 2). The moderate correlations between DSB across cell types demonstrate that a substantial proportion of DSB susceptibility across the genome is cell type specific, which is consistent with the established cell type specific properties of many SV breakpoint regions in tumours, such as common fragile sites (38). Furthermore the larger performance gap in models for cell lines with altered variable rankings indicates that DSB mechanisms may differ across cell types and may not be completely captured via epigenomic features.

**Figure 4:**
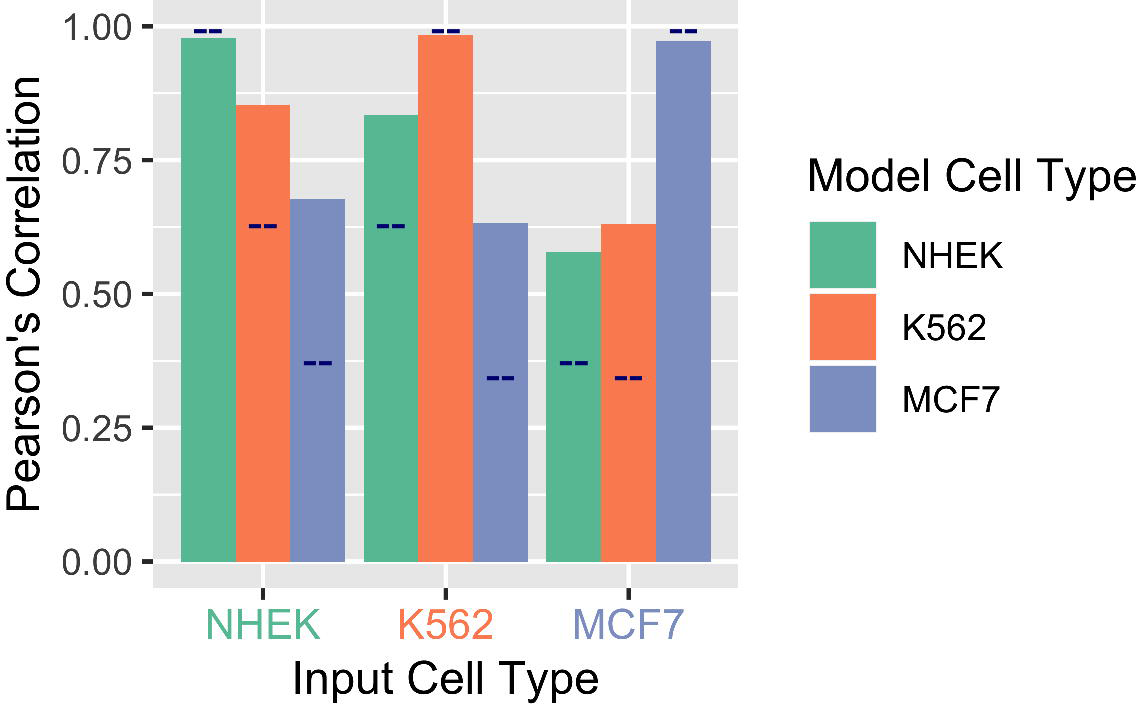
DSB models improve predictions for non-model cell types. Models trained using a dataset from one cell type were used to generate predictions for a different cell type, given the matched features. The dark blue lines mark the Pearson′s correlation between the two cell types. The cell type used to train the model is indicated by the colour of the bar, and the cell type on which the model is being applied is shown on the x-axis. In all cases, the random forest model greatly improves the predictions from a naive inference, with a 1.3-1.8 fold improvement in correlation.

## Tumour SV breakpoints possess variable susceptibility to DSBs

Keratinocytes are considered to be the cell type of origin for mucosal and cutaneous carcinomas, particularly squamous cell carcinomas (39), and NHEK cells are often used in the literature as a model for these cancers. Similarly, MCF7 cells and K562 cells have been used extensively as models for breast and blood cancers respectively. This motivated us to ask how the DSB models for these three cell types relate to the patterns of SV breakpoints observed in squamous cell carcinomas, blood cancers, and breast tumours.

A number of large structural variant (SV) collections have been established for a variety of tumour types, and each possesses advantages and shortcomings. The International Cancer Genome Consortium (ICGC) provides high resolution SV calls based upon whole genome sequencing (WGS) for 2,146 patients across 17 cohorts (40), but sample cellularities, sequencing depths and SV calling methods vary across cancer cohorts, and are expected to affect results (Supp Figure 6). The Cancer Genome Atlas (TCGA) produced consistently processed copy number variant (CNV) calls from SNP chip data for 23,084 patients across 33 cohorts (Supp Figure 7). However, breakpoint resolution is much lower than calls based upon WGS, and copy neutral SVs such as inversions and translocations are absent. We analyzed ICGC and TCGA data as pancancer datasets, combining all cancer types together, but also as three cancer type subgroups. TCGA subgroups comprised a squamous cell carcinoma subgroup, a blood cancers subgroup including two blood cancers, and breast cancer as a separate group (see Methods). Similar ICGC subgroups were formed (from cohorts independent of TCGA), but with the squamous cell carcinoma subgroup replaced with a carcinoma subgroup, which includes seven carcinoma cancer studies excluding breast cancer (see Methods).

Analogously to the DSB datasets, we determined the number of tumour SV breakpoints per 50kb region for each of the ICGC and TCGA SV datasets (see methods) and compared these to the DSB predictions from our models. In ICGC data overall we saw low correlations between the number of SV breakpoints and DSB predictions (Supp Figure 8 and Supp Figure 9). Restricting our analysis to ICGC enriched SV breakpoint regions, or ESBs for the purpose of this manuscript (50kb regions with SV breakpoint counts in the top 5% genome-wide, see Methods), increased the agreement with DSB model predictions. Significant increases in NHEK and MCF7 model predictions were seen for pancancer, carcinoma, blood, and breast tumour ESBs and in K562 model predictions for all cancer subsets except blood ESBs (Figure 5). The significant increase in DSB model predictions seen for carcinoma ESBs indicates that DSB susceptibility (captured in the models) may shape the SV landscape of these cancer types. We also see a significant increase in DSB predictions for TCGA blood cancer ESBs, but not for any other subgroups in TCGA data (Supp Figure 10). However, as mentioned, TCGA data is of low resolution and not suitable for accurate breakpoint detection.

**Figure 5:**
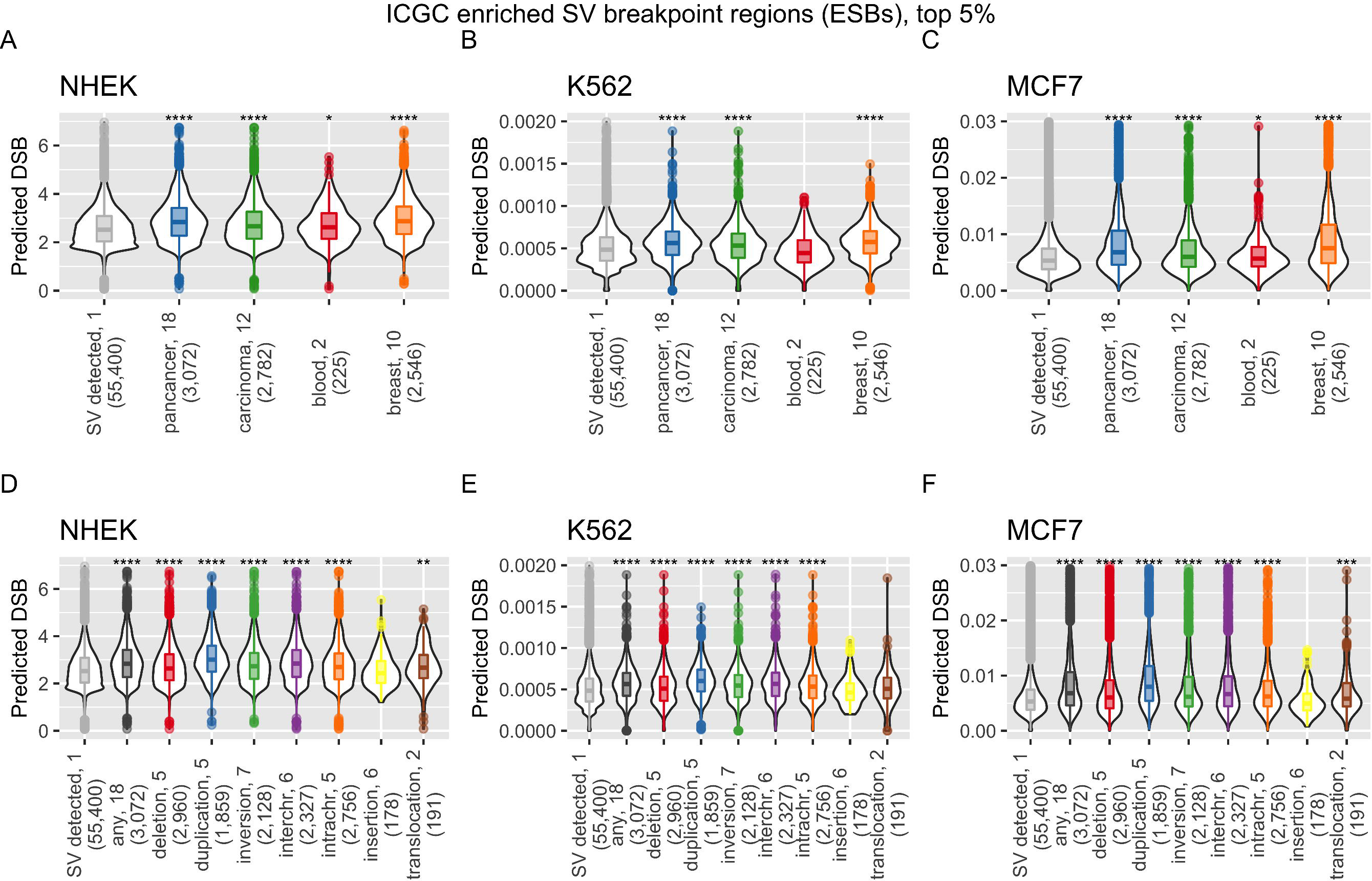
Regions enriched for cancer SV breakpoints (ESBs) display a significant increase in DSB frequency across cancer types. A-C) The regions with ICGC SV breakpoint frequencies in the top 5% are shown with their predicted DSB values as violin plots for each of the three cell type models: NHEK, K562, and MCF7. ICGC cohorts are shown all together (pancancer), and split into three cancer categories: carcinoma, blood, and breast cancers (see methods). D-F) ICGC SV breakpoint counts separated by SV type, and the top 5% of ESBs are shown with their predicted DSB values as violin plots. The numbers following the x-axis labels are SV breakpoint count cut-offs for the top 5% ESBs, and the numbers in parenthesis are the number of 50kb regions that meet the cut-off. For example, there are 225 50kb regions with more than two SV breakpoint in blood cancers. Stars indicate significantly higher values in DSB predictions for the ESBs relative to non-ESBs for each category, as determined by a Wilcox ranked sum test (* for p<=0.05, ** for p<=0.01, *** for p<=1e-3, and **** for p<=1e-4).

Certain classes of relatively simple SVs (deletions, duplications, inversions, translocations) are often the product of one or two DSBs, while more complex intrachromosomal rearrangements can be difficult to classify accurately, and may have origins in poorly understood phenomena such as chromothripsis (41). Indeed, even for simple SVs there may be some ambiguity, with an unknown fraction arising by mechanisms that may not involve a DSB. For example, insertions can arise from transposon activity, and duplications from replication slippage (42). However, even if many SV breakpoints do not arise from DSBs, we might reasonably expect to see shifts to higher median DSB model prediction values for many simple SV classes. We determined ESBs as above for ICGC-annotated SV classes across all ICGC tumour types to examine their DSB frequency predictions, compared to non-ESBs, 50kb regions that do not attain SV breakpoint counts in the top 5% with at least one tumour SV breakpoint detected. Overall, the models show significant elevations for ESBs covering all SV classes except insertions (Figure 5). Insertions may be less influenced by DSB susceptibility because they may occur via transposable element activity rather than through DNA damage and repair pathways. Crosetto et al. (18) find an enrichment of satellite repetitive elements in regions enriched for DSB in cells exposed to aphidicolin. However, regions that undergo DSB under replicative stress, as induced by aphidicolin, may differ from DSB regions under normal cell growth conditions.

## Interrogating tumour SV data at common fragile sites with DSB models

The predicted DSB frequencies from our models and ICGC tumour SV breakpoint frequencies differ in their scaling and distributions and are not directly comparable. However, it is of interest to identify outlier regions, where model predictions and observed tumour SV breakpoint rates diverge most, since these regions may include loci under selection in tumours. We developed a novel metric, the d-score, to measure this divergence between expectations given a DSB model and observed SV breakpoint rates in tumours. In brief, this metric relies on fitting known distributions to the observed SV breakpoint dataset and to the predicted DSB dataset. Based upon the known distributions we then transform the observed SV counts and predicted DSB values to p-values, reflecting the probability that each value is drawn from the fitted distribution (see Methods). For each 50kb region in the genome the difference between the SV breakpoint log p-value and the predicted DSB log p-value is the d-score. Regions with unexpectedly high d-scores contain more SV breakpoints than expected, given our model, whereas regions with unusually low d-scores contain fewer SV breakpoints than expected.

Common fragile sites (CFSs) have long been studied for their unusual properties of generating SVs, both in normal cells and in cancer (38). These regions undergo frequent DSBs in tumours and have been well studied in terms of their genomic context, relationship to replication timing and origins, and correlations with particular chromatin states (43). They tend to occur within large genes, in G-negative chromosomal bands with high DNA flexibility, are unusually late replicating (44), and it is thought that their instability derives from transcription-associated replication stress (38). CFSs only exist in modest numbers and are defined at low resolution (by cytogenetic bands or gene loci); they therefore provide an interesting, though challenging, test set of regions to examine d-score performance.

We examined predicted (NHEK model) DSB frequencies at 294 50kb regions coinciding with annotated CFS gene loci across the genome, in comparison to regions associated with all annotated genes, and regions associated with putative cancer driver genes (Figure 6C). Although significant shifts to higher frequencies are seen for the driver gene sets for predicted DSB frequencies, the CFSs do not show a similar increase, most likely because the model predicts DSB in early replicating regions, and CFS tend to be late-replicating. Thus, the dominant features influencing DSB susceptibility genome-wide do not appear to drive the elevated DSB rates at CFSs, consistent with CFS instability involving replicative stress (38). However, CFS d-scores show a significant shift above the distribution for all genes and above the driver gene sets as well (Figure 6D). This result is replicated in the MCF7 BLISS model examined inconjunction with ICGC breast cancer SV breakpoints (Sup Figure 11). We conclude that the d-score, a measure of relative DSB enrichment, offers a robust metric for the classification of regions showing unusual SV breakpoint rates in tumours.

**Figure 6:**
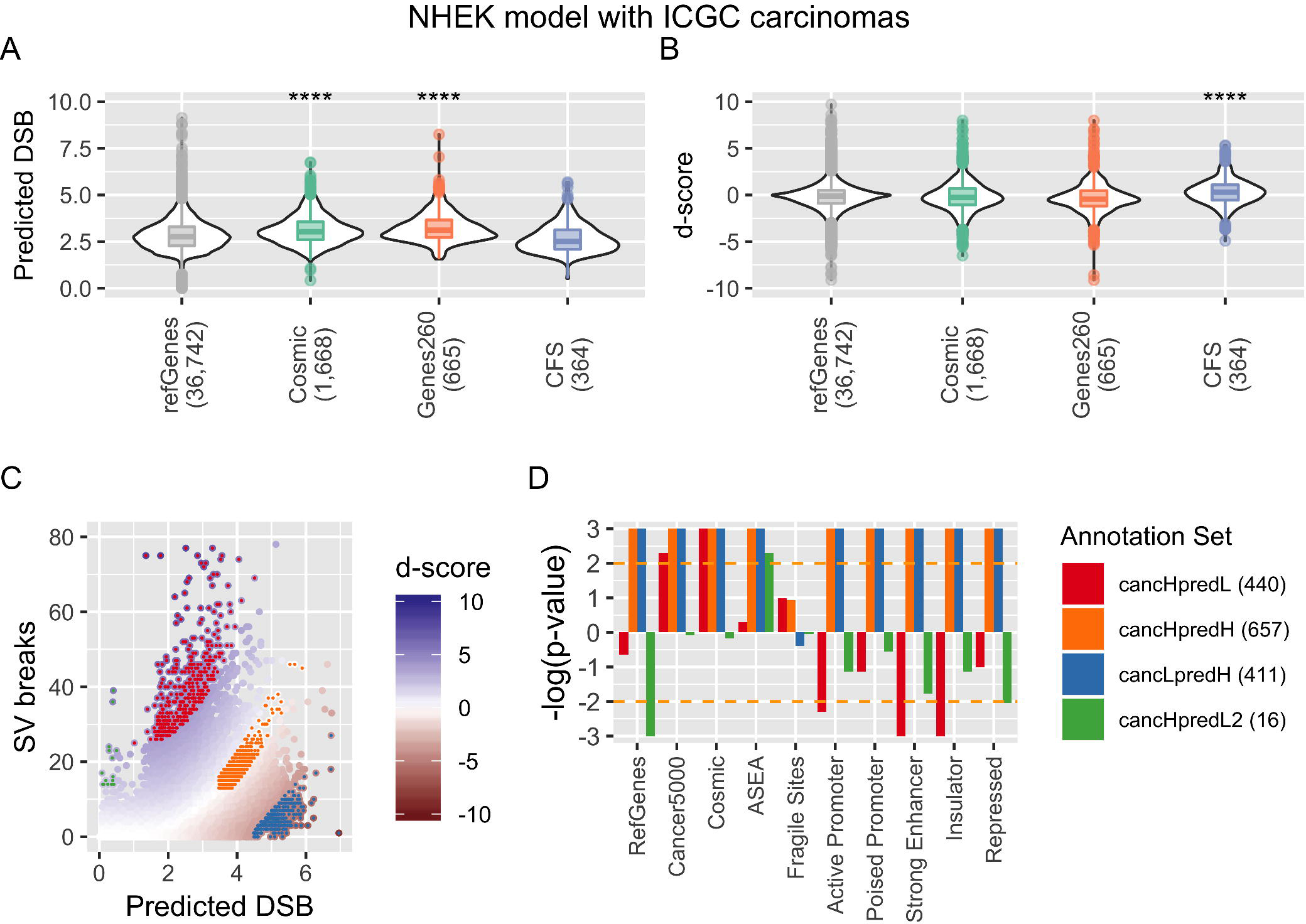
Inference of positively and negatively selected SV regions. A) The predicted DSB frequencies for regions overlapping RefSeq genes, two sets of cancer consensus genes, and common fragile sites (CFS) are shown as violin plots. The stars represent significantly higher values in the region subsets, compared to genomic regions that do not overlap the given annotation set, using a Wilcox ranked sum text. B) The same regions as in a), but with d-score values, a measure of the deviation of the observed breakpoint frequencies from the predicted or expected DSB frequencies. C) Observed SV breakpoint frequencies for ICGC carcinomas (excluding breast cancer) with predicted DSB frequencies from the NHEK DSBCapture model. Each point represents a 50kb region and is coloured by its d-score. Regions were split into high (cancHpredL) and low (cancLpredH) d-score categories (d-score p-value<0.01), a cancHpredH category, representing regions with d-scores near zero, and a cancHpredL2 category, representing low mappability regions (see methods). D) Each category was tested for enrichment of various annotations using circular permutation (see methods). The yellow dotted line marks p<0.01 significance, and the numbers in parenthesis indicate the number of 50kb regions in each category, out of 61,903 in total.

## Identification of hot and cold spots for structural variant breakpoints in tumours

We have developed a classification of regions of interest within ICGC tumour cohorts based upon the d-score metric. We call regions with significantly more SV breakpoints than expected, or SV hotspots, cancHpredL (cancer high, predicted low), and regions with fewer SV breakpoints than expected, or SV coldspots, cancLpredH (cancer low, predicted high) (see Methods). Figure 6 depicts these classes of regions in d-score plots of ICGC SV breakpoint data. Many previous studies have predicted oncogenic SV hotspots simply as regions repeatedly rearranged in cancers. Here we refine such predictions by assessing these raw SV breakpoint frequencies relative to the predicted susceptibility of each region to breakage. It is not possible to predict coldspot regions without a model of expected DSB frequency, and to our knowledge SV breakpoint coldspots have not been studied before.

We also define a class of regions possessing both high predicted DSB values and high SV breakpoint frequencies (cancHpredH), corresponding to regions showing unusually high SV frequencies on the background of high susceptibility to DSBs. Finally, we define a fourth class of regions that have predicted DSB rates close to zero but high SV breakpoint frequencies (cancHpredL2). In principle, these regions are a class of SV hotspots but, as shown in Figure 3B, they are likely to be repetitive, heterochromatic, and enriched for artifacts (false positives and negatives in SV breakpoint) due to their association with low mappability.

We examined a range of functional annotation enrichments in the four classes of regions using circular permutation to assess significance (see Methods; Figure 6). The annotations included two putative cancer gene sets, 260 genes from the Cancer5000 dataset (45) and 561 genes from the COSMIC collection (46)). We also included a set of 15,415 super enhancers (47), common fragile sites, and chromatin states from ENCODE chromHMM analysis (48). Notably, the majority of genes in both cancer sets are predicted to be oncogenic based on unexpectedly high and functionally significant SNV (rather than SV) loads and are not necessarily expected to occupy regions with higher levels of SV breakpoints. In fact, both gene sets demonstrate significant enrichments in the cancHpredL class of hotspot regions (Figure 6D), although RefSeq genes do not, suggesting that these genes may also frequently be altered in cancer through SV. The cancHpredL regions are also significantly depleted in active chromatin regions, such as promoters, enchancers, and insulator regions, most likely because these types of regions do not have low predicted DSB. The high susceptibility cancHpredH regions occupy gene-rich areas of the genome (enriched for known RefSeq genes) including both cancer genes sets, and for active promoters, strong enhancers, and insulators. This is consistent with reports that CTCF bound insulator elements suffer recurrent mutations in tumours. Likewise, the cancLpredH class of coldspot regions occupy gene rich neighbourhoods, active promoters, and strong enhancers (Figure 6), suggesting some genes and distal regulatory regions may have experienced purifying selection in tumours.

Given the discrepancies mentioned above between ICGC and TCGA experimental platforms, data analysis, and sample cohorts, we do not expect strong agreement between ICGC and TCGA derived SV datasets. Indeed, the correlation between them is low (Spearman’s rho of 0.099, p<2.2e-16), and the pancancer ESBs from either set do not significantly overlap (p < 0.99, see methods). However, the cancLpredH class is again enriched in active promoter and strong enhancer regions, in accordance with the results based upon ICGC SV data (Sup Figure 12).

We again wanted to test the utility of DSB random forest models applied to different cell types by testing the accuracy of predictions made by a model trained in one cell type given features for a different cell type, as in Figure 4. Instead of looking at the correlation between the observed and predicted DSB scores across the genome, we examined the overlap between cancHpredL, cancHpredH, and cancLpredH 50kb regions for the MCF7 model versus the NHEK model, using the MCF7 model as the truth set. Subsets of 50kb regions for each model were derived from MCF7 features and ICGC breast cancer SV breakpoints; only the training data for the models differ. We found a significant overlaps between all three categories of d-score subsets, with 595/662 cancHpredL, 255/785 cancHpredH, and 253/594 cancLpredH regions detected via the NHEK model (p<2.2e-16), demonstrating that a given model can be used to detect regions of interest in various cell types.

## Functional annotation of regions of interest

We closely examined the ten 50kb regions with the highest (cancHpredL) d-scores to uncover genes that might be reclassified as oncogenic due to a higher than expected SV breakpoint frequency in cancer. Likewise, we investigated the ten regions with the lowest d-scores (cancLpredH), which we predict to be under purifying selection, for signals of potential functionality. For this analysis we used the NHEK model predictions paired with ICGC carcinoma SV breakpoints.

Nine out of ten regions with the highest d-scores overlap a gene, and four overlap COSMIC genes. *CHEK2* and *CDKN2A* are known tumor suppressors, and *TMPRSS2* and *ERG* is frequently involved in translocation events forming fusion oncogenes in certain cancers. For example, it fuses with *TMPRSS2* in most prostate cancers, with *EWS* in Ewing’s sarcoma, and with *FUS* in AML. Two adjacent 50kb regions on *chr17q12* overlap *GRB7* and *IKZF3*. *GRB7* encodes a protein that interacts with epidermal growth factor receptor *(EGFR)*, a well-known proto-oncogene, and *IKZF3* is a zinc finger protein and transcription factor involved in B lymphocyte regulation and differentiation as well as chromatin remodeling. This region also corresponds to a known fragile site *FRA17A* (49). Of the ten regions with the lowest d-scores, seven overlap a known gene and two known oncogenes. The oncogene, *CDC27*, or cell division cycle 27, encodes a component of the *APC* and has been shown to interact with other mitotic checkpoint proteins. It is highly conserved and may be necessary for cell survival. There is also a non-coding RNA found on chr2 in the centromeric region, *LOC654342*, which overlaps an H3K27ac peak, and may be acting as a regulatory element.

## Discussion

Recent *in vitro* studies of DSB frequency in cell lines have suggested that a variety of underlying genomic features are associated with DSB susceptibility. We have shown that accurate models of genome-wide DSB frequency can be built from a modest number of such features, with replication timing, open chromatin, and marks of active promoter or enhancer regions associated with increased DSBs. Although active regulatory regions often harbor actively transcribed genes, it appears that chromatin accessibility at these sites rather than transcription itself determines DSB propensity. The variable importance metrics also show certain features to be more influential in particular cell types, with CTCF and H3K36me3 having more predictive power in MCF7 than in NHEK or K562. Not only are DSB patterns cell type specific, but the factors influencing those patterns also depend on cell type, suggesting different mutational mechanisms at play. As a matter of course, our models’ accuracies decline when applied to cell lines other than the training set, but they still generate reasonable DSB frequency predictions, with correlations between 0.57 and 0.83 to the observed data, which are large improvements over a simple inference. Since chromatin features influence mutation patterns and are cell type specific, it will be important to use mutational propensity profiles for matched cell types in future cancer studies.

Our models of genome-wide DSB susceptibility predict DSB frequencies for all 50kb loci, and reflect the established correlations between replication timing and DSB frequency (50) as well as tumour SV rates (9,10). A recent complementary study has shown that 84,946 high confidence peaks of NHEK DSBCapture signal (22), marking small (median: 391bp) sites of unusually high DSB susceptibility, can be accurately classified from control sites using underlying genomic features (34). Consistent with our results, this binary classifier suggested prominent roles for DNase accessible regulatory sites and CTCF binding, and recapitulated many of the patterns reported by Lensing et al (2016). However, the model of Mourad et al (2018) omitted replication timing and does not provide quantitative predictions of DSB susceptibility across the genome.

We used our genome-wide models of DSB susceptibility to interrogate the largest tumour SV breakpoint collections and found surprising levels of agreement, such that SV breakpoint enriched regions often show shifts to higher predicted DSB susceptibility. In spite of variable sample sizes, the classes of simple SV likely to arise by one or two DSBs (deletions, duplications, inversions, translocations) showed significant increases in predicted DSB susceptibility. The NHEK model best predicted the patterns of DSB susceptibility in tumours, showing genome-wide elevations of predicted DSBs for all of these SV classes relative to control regions. Thus, the chromatin-mediated DSB susceptibility captured in the model may shape the landscape of SV recurrence in these classes.

There are many reasons why one might expect a much poorer agreement between the predictions of in vitro DSB frequency models and the patterns of SV breakpoints observed in tumour sequencing studies. The available collections of SV breakpoints in tumours are far from perfect, and even the best ICGC data suffer large variations in sample size, sample heterogeneity, sequencing depths and SV calling methods across tumour cohorts. In addition, fundamental aspects of tumour biology (cellular heterogeneity, disrupted repair pathways, chromatin alterations etc.) are expected to place distinct limits on the agreement we can see with the DSB patterns seen in cell lines. Evidence is also emerging that there are important properties of the mutational landscape in tumours that are unlikely to be captured by in vitro model systems. For example, a recent study of intra-tumour diversification in colorectal cancer suggests that most mutations occur during the final clonal expansion of these tumours, resulting from mutational processes that are absent from normal colorectal cells (51). Enhanced rates of DSB formation have also been observed in vitro at cryptic replication origins activated by oncogene-induced replication stress, though these cryptic sites seem to explain only a minority of SV breakpoints (<8%) across a variety of TCGA tumour types (52). Given the many known and possible differences between in vitro DSB model predictions and observed tumour SV breakpoints, it is remarkable that significant agreement is found on any level.

There is great interest in ‘hotspot’ genomic regions harbouring recurrent SVs in tumours, on the basis that such regions may be under positive selection, conferring a proliferative or survival advantage to tumour cells. However, rigorous inference of selection requires a proxy for the expected rate of recurrence within such regions. Using model predictions as this proxy we have produced refined hotspot predictions, reflecting SV breakpoint frequencies relative to the predicted susceptibility of each region. Since our predictions of DSB susceptibility are genome-wide it was also possible to predict coldspot regions, regions possessing unexpectedly low SV breakpoint rates given model predictions, and putatively subject to negative or purifying selection in tumours. If selection in tumours is prominent in driving SV breakpoint frequencies away from DSB model predictions, we might expect hotspot and coldspot regions to show unusual functional enrichments. Multiple caveats apply to the annotations examined but analysis using the NHEK model shows that ICGC carcinoma hotspots are enriched for putative oncogenes. Coldspots occupy gene-rich neighbourhoods but and are also enriched in active promoters and strong enhancers, and insulators, indicating regulatory regions that may have experienced purifying selection in tumours.

## Conclusions

When inferring selection on single nucleotide variants it is standard practice to make comparisons between the observed variant frequencies and the frequencies expected, according to a model of single nucleotide mutation rates. We have developed models of DSB mutation rates that can be used to generate expected SV breakpoint frequencies and illuminate regions with significant deviations from these expectations. This approach provides statistically rigorous protocols to prioritize novel loci putatively under selection in tumours, generating testable hypotheses for further experimental studies.

## Methods

### Derivation of DSB data in the K562 and MCF7 cell lines

DSB profiles were generated with an adapted version of the Breaks labeling *in situ* and sequencing protocol (25), in which DSB ends are labeled with a dsDNA BLISS adapter in cell suspensions of 1 million cells. Afterwards the published protocol is followed with only minor modifications. Labeled DSBs are selectively amplified using T7-driven linear amplification, after which sequencing libraries are generated and sequenced with single-end 1×75 v2 chemistry on an Illumina NextSeq 500. Raw sequencing reads were demultiplexed by Illumina’s BaseSpace, after which FASTQ files were downloaded and processed as described in Yan et al. 2017 (SRA accession SRP150602). In brief, reads with the expected prefix of 8nt UMI and 8nt sample barcode sequence were filtered using SAMtools and *scan for matches,* allowing at most one mismatch per barcode. Trimmed reads were then aligned to GRCh37 using bwa mem, and reads with mapping scores below 30 were discarded. Next, PCR duplicates were identified by searching for proximal reads (within 30bp of the reference genome) with at most two mismatches in the UMI sequence, which were then grouped and collapsed into a single break location. Finally, we generated .bed files with DSB locations and the number of unique UMIs indicating that location.

### Generating random forest models

We downloaded ten tracks from ENCODE for multiple chromatin marks, replication timing, open chromatin, several DNA binding proteins, and nucleosome pull-downs from the UCSC genome browser (53). We used G-quadruplex data generated by Chambers et al, (GSE63874). In their study, they make separate .bedgraph files available with the G-quadruplex density for each strand. We used the sum of the plus and minus strands in our analysis. The list of bigwig files used for each cell line along with their sources and graphical labels is in Supplementary Table 1. We used the bigWigAverageOverBed tool from the kentUtils tool library to produce average signal per 50kb in non-overlapping windows across hg19 for each track. We combined the results to a single matrix per cell line composed of 61,903 rows, one for each 50kb bin, and 11 columns, one for each chromatin or genomic feature. These feature matrices are available in supplementary data and scatter plots of each feature with the NHEK DSBCapture data are shown in Supplementary Figure 3.

For the extended model in Supplementary Figure 4, we downloaded an additional nine features from the UCSC genome browser (53), which were processed in the same way as the ten ENCODE features used in the primary feature matrix. We also downloaded .hic files for NHEK, K562, and HMEC cells generated from Rao, et al. (GSE63525). We used their custom toolbox, Juicer, to calculate eigenvectors per chromosome, and generated 50kb resolution eigenvector profiles using the bedGraphToBigWig and bigWigAverageOverBed tools from kentUtils. The figure labels and sources for these data are in Supplementary Table2, and the extended feature matrices are in supplementary data.

We generated DSB frequency scores from each of four HTS DSB profiling datasets: two in NHEK cells, one for K562, unpublished, and one for MCF7, unpublished. As mentioned in the results, two replicates for each of two DSB HTS profiling methods, DSBCapture and BLESS, were available from Lensing et al. (22). We took the average per 50kb of the replicates to create an NHEK DSBCapture profile and an NHEK BLESS profile. We combined three replicates of MCF7 BLISS data (via a sum operation) to serve as our MCF7 DSB profile. A fourth MCF7 BLISS dataset is available, but we excluded it from our analysis because it had a distinctly lower correlation to the other three datasets (0.90-0.92 as opposed to 0.97-0.99). These scores are available as supplementary files.

We used the randomForest package in R to generate random forest models with 500 trees and five OOB permutations per tree (options ntree=500, nPerm=5). To calculate variable importance, we used the importance command within the randomForest package (https://cran.r-project.org/web/packages/randomForest/index.html), which calculates the average prediction error rate (MSE) for each datapoint (50kb bin) across all trees in the random forest. Then, for each feature variable, the values are randomly permuted and the MSE for each 50kb bin is calculated again. The final variable importance score is the average difference in MSE before and after the permutation, normalized by the standard deviation of these differences. Because many features are inter-correlated, their importance measures were very similar. Therefore, in order to determine a consistent ranking of features’ importance values, we generated ten random forest models per dataset and calculated the average and standard deviation of importance across the ten models.

Although random forest models are not susceptible to overfitting, to confirm that our models were not overfit to the DSB data, we also generated a random forest model for the NHEK DSBCapture dataset, holding out one third of the data as the test set and training the model on the remaining two thirds. This model showed 0.93 Pearson’s correlation between the predictions and the observed data for the training set, similar to the model trained on the full dataset (Sup Figure 5).

### Determining tumour ESBs and their predicted DSB scores

To determine SV DSB rates in from TCGA data, we downloaded CNV data from TCGA (54), which came from Affymetrix SNP 6.0 arrays processed by the DNAcopy R-package (https://docs.gdc.cancer.gov/Data/PDF/DataUG.pdf). DNAcopy generates a set of continuous segments, outputting regions with little or no copy number change, so we filtered these, defining segments with a CN ratio >1 as amplifications and ratios < −1 as deletions. The segments were lifted from hg38 to hg19 using UCSC’s liftOver tool. For each CNV, we counted a single DSB to occur in a 50kb bin if either or both ends of the segment overlapped the bin. The TCGA-BLOOD group includes the two blood cancer cohorts: acute myeloid leukemia (LAML) and lymphoid neoplasm diffuse large B-cell lymphoma (DLBC), while the TCGA-SCCA group includes three squamous cell carcinomas: cervical squamous cell carcinoma and endocervical adenocarcinoma (CESC), head and neck squamous cell carcinoma (HNSC), and lung squamous cell carcinoma (LUSC). The BRCA group includes only the TCGA breast cancer cohort (BRCA), and the PANC group includes all 33 cancer types, shown in Supplementary Figure 7. Counts for various groups and CNV types are available as Supplementary Files.

We downloaded available WGS SV calls from the ICGC Data Portal (https://dcc.icgc.org/projects). As with the TCGA CNV, a single DSB was counted per 50kb bin if either one or two ends of a SV overlapped the region. The ICGC pancancer group contains SVs from 17 cancer studies, shown in Supplementary Figure 6. The carcinoma group contains all available carcinoma cancer studies, excluding breast cancer: early onset prostate cancer (EOPC-DE), liver cancer (LIRI-JP), pancreatic cancer (PACA-CA, PAEN-AU, PAEN-IT), prostate cancer (PRAD-CA, PRAD-UK), and skin adenocarcinoma (SKCA-BR). The ICGC blood group contains chronic lymphocytic leukemia (CLLE-ES) and malignant lymphoma (MALY-DE), and the breast group contains breast cancer studies (BRCA-EU and BRCA-FR). A table of DSB counts per 50kb broken up by group and SV type is in supplementary data.

We determined enriched SV breakpoint regions (ESBs) per cohort or SV type grouping by ranking the 50kb bins by the number of DSB, excluding regions with no DSB in the group, and using the number of DSB in the top 5% as the cutoff. All 50kb regions with a DSB count greater than or equal to the cutoff were designated ESBs. We used a Wilcoxon ranked sum test (R wilcox.test command) to test for significant increase in the predicted DSB values for ESBs compared to all other regions, and we excluded regions in which no DSB were found in any cancer study since these are likely to be unmappable or blacklisted regions.

The correlation between TCGA and ICGC pancancer SV breakpoint counts was calculated using Spearman’s rho and excluding 50kb regions with no SV breakpoints in either the TCGA or ICGC datasets. The top 5% ESBs were found for each dataset, with 2,839 regions found in TCGA and 3,072 in ICGC, and the significance of the overlap was calculated using a hypergeometric test (R command phyper with q=177, m=2,839, n=61,903-2,839, and k=3,072).

### Calculating d-scores

We used the R package fistdistrplus (55) to determine the distributions with the best fit to the DSB prediction values and the SV breakpoint frequencies. We used a likelihood maximization test (method=“mle”) and the BIC (Bayesian Information Criterion) measure of goodness of fit to choose the best distribution. We tested a lognormal, log-logistic, gamma, normal, and an exponential distribution, and fitted the distributions to the bulk of the SV breakpoint or DSB prediction data. We excluded 50kb regions with breakpoint frequencies greater than six times the interquartile range from the median in order to exclude extreme outliers. While we aimed to emphasize the fit of the tails of our data’s distributions, including these outliers resulted in poorly fitting distributions to the bulk of the real data. Once we found the best of the three candidate model distributions, we assigned a p-value to each 50kb bin from the fitted distribution (using the plnorm, pllogis, or pgamma functions in R) which represent the probability of seeing a given breakpoint frequency or DSB prediction or greater in the known distribution. The actual and fitted distributions and quantile-quantile plots are shown in Supplementary Figures 13 and 14.

Next, for each 50kb bin, we calculated the difference in log p-values between the predicted DSB and the actual SV breakpoints, called d-scores. Using the fistdistrplus R package again, we determined the best-fit distribution for the d-scores, choosing between a t-distribution, a normal, and a Cauchy distribution. Again, we used a maximum likelihood method and the BIC measurement and excluded extreme outliers. In all cases, a t-distribution with four degrees of freedom (df=4) was the best fit, so each 50kb bin was assigned a p-value from this distribution according to its d-score. The histograms and quantile-quantile plots of the d-scores and fitted distributions are shown in Supplementary Figure 15.

### Calculating gene set and chromatin domain enrichments

We used the d-score p-values to categorize regions into informative subsets, using the R command qt(p=0.01, df=4, lower.tail=FALSE) to determine the d-score cutoffs. The cancHpredL class of regions have d-scores in the upper one percentile (> 3.75), and the cancLpredH have d-scores in the lower one percentile (< −3.75). The cancHpredH class has d-scores in the 40^th^ to 70^th^ percentiles and SV breakpoint frequencies or DSB predictions with p-values less than 0.01, so these regions have significantly (p-value < 0.01) high SV breakpoints or DSB predictions but insignificant d-scores (p-value < 0.6). The cancHpredL2 class consists of regions with SV breakpoint p-values less than 0.01, and DSB predictions less than 0.5 for the NHEK models and less than 0.001 for the MCF7 model.

We used a binomial test to measure the significance of overlaps between sets when comparing results from the MCF7 model and the NHEK model applied to ICGC breast cancer data and MCF7 cell line features (R command binom.test).

We used the R package regioneR (56) to compute the overlap significance between each set of regions and various genome and chromatin annotation files. A list of annotation sets and their original sources are in Supplementary Table 2. We matched Cancer5000 genes and Cosmic gene lists to RefSeq gene names in order to get their genome coordinates, so the cancer gene lists are RefSeq gene subsets. The super enhancer set (SEA) came from A549 cells, derived from a lung carcinoma (47). Common fragile sites (CFS) were collected from NCBI’s gene archive by searching for “common fragile site” or “fragile site” within human genes. Many fragile sites are annotated by chromosome band but do not have exact coordinates; we filtered these out because they are low resolution. The chromHMM (48) annotation came from the UCSC genome browser. We tested enrichment of the NHEK states with the NHEK model d-score classes and the HMEC track, from primary mammary epithelial cells, with the MCF7 model’s d-score classes. The regioneR package performs random circular permutation of regions of interest and then computes the number of overlaps between the permutated set and a second set of regions. The p-value represents how often, over the course of the permutations, the two sets overlap to the same extent that they do without any permutation. We used 1,000 iterations to achieve a maximum p-value of 0.001.

## Declarations

### Ethics Approval

Approval for access and use of ICGC variant data was obtained from the ICGC Data Access Compliance Office. Use of TCGA CNV does not require ethics approval.

### Consent for Publication

Not applicable

### Availability of data and materials

All analysis was done using GRCh37 as the reference genome. The raw BLISS sequencing data is available on SRA with accession SRP150602. All scripts and commands used to do this analysis are available on github (https://github.com/TracyBallinger/dsb_model). In addition, we have made ipython notebooks for the figures used in this manuscript to ease reproducibility and allow further exploration of the data, also available on github. All supplementary files are available for download at https://datashare.is.ed.ac.uk/handle/10283/3103.

## Authors’ Contributions

BB and RM generated the BLISS DSB profiles. SG developed the BLISS alignment pipeline and generated .bed files of DSB profiles. TB performed all subsequent data analysis and produced figures. TB and CS wrote the manuscript. NC and CS supervised the project. TB, BB, NC, and CS edited the final manuscript.

## Acknowledgements

We are indebted to the ICGC and TCGA projects for the timely public release of tumour genome sequencing data and SV calls.

## References

1. Ciriello G, Miller ML, Aksoy BA, Senbabaoglu Y, Schultz N, Sander C. Emerging landscape of oncogenic signatures across human cancers. Nat Genet. 2013 Oct;45(10):1127–1133.

2. Patch A-M, Christie EL, Etemadmoghadam D, Garsed DW, George J, Fereday S, et al. Whole-genome characterization of chemoresistant ovarian cancer. Nature. 2015 May;521(7553):489–494.

3. Scarpa A, Chang DK, Nones K, Corbo V, Patch A-M, Bailey P, et al. Whole-genome landscape of pancreatic neuroendocrine tumours. Nature. 2017 Mar;543(7643):65–71.

4. Alaei-Mahabadi B, Bhadury J, Karlsson JW, Nilsson JA, Larsson E. Global analysis of somatic structural genomic alterations and their impact on gene expression in diverse human cancers. Proc Natl Acad Sci U S A. 2016 Nov;113(48):13768–13773.

5. Li Y, Roberts N, Weischenfeldt J, Wala JA, Shapira O, Schumacher S, et al. Patterns of structural variation in human cancer. bioRxiv. 2017 Aug;181339.

6. Sudmant PH, Rausch T, Gardner EJ, Handsaker RE, Abyzov A, Huddleston J, et al. An integrated map of structural variation in 2,504 human genomes. Nature. 2015 Oct;526(7571):75–81.

7. Weischenfeldt J, Dubash T, Drainas AP, Mardin BR, Chen Y, Stütz AM, et al. Pan-cancer analysis of somatic copy-number alterations implicates IRS4 and IGF2 in enhancer hijacking. Nat Genet. 2017 Jan;49(1):65–74.

8. Glodzik D, Morganella S, Davies H, Simpson PT, Li Y, Zou X, et al. A somatic-mutational process recurrently duplicates germline susceptibility loci and tissue-specific super-enhancers in breast cancers. Nat Genet. 2017 Jan;49(3):341–348.

9. Morganella S, Alexandrov LB, Glodzik D, Zou X, Davies H, Staaf J, et al. The topography of mutational processes in breast cancer genomes. Nat Commun. 2016 May;7:11383.

10. Schuster-Böckler B, Lehner B. Chromatin organization is a major influence on regional mutation rates in human cancer cells. Nature. 2012 Aug;488(7412):504–507.

11. Ding L, Getz G, Wheeler DA, Mardis ER, McLellan MD, Cibulskis K, et al. Somatic mutations affect key pathways in lung adenocarcinoma. Nature. 2008 Oct;455(7216):1069–1075.

12. Jackson SP, Bartek J. The DNA-damage response in human biology and disease. Nature. 2009 Oct;461(7267):1071–1078.

13. Biehs R, Steinlage M, Barton O, Juhász S, Künzel J, Spies J, et al. DNA Double-Strand Break Resection Occurs during Non-homologous End Joining in G1 but Is Distinct from Resection during Homologous Recombination. Mol Cell. 2017 Feb;65(4):671–684.e5.

14. Nussenzweig A, Nussenzweig MC. A backup DNA repair pathway moves to the forefront. Cell. 2007 Oct;131(2):223–225.

15. Clouaire T, Legube G. DNA double strand break repair pathway choice: a chromatin based decision? Nucl Austin Tex. 2015;6(2):107–113.

16. Glover TW, Berger C, Coyle J, Echo B. DNA polymerase alpha inhibition by aphidicolin induces gaps and breaks at common fragile sites in human chromosomes. Hum Genet. 1984;67(2):136–142.

17. Canela A, Sridharan S, Sciascia N, Tubbs A, Meltzer P, Sleckman BP, et al. DNA Breaks and End Resection Measured Genome-wide by End Sequencing. Mol Cell. 2016 Sep;63(5):898–911.

18. Crosetto N, Mitra A, Silva MJ, Bienko M, Dojer N, Wang Q, et al. Nucleotide-resolution DNA double-strand break mapping by next-generation sequencing. Nat Methods. 2013 Mar;10(4):361–365.

19. Frock RL, Hu J, Meyers RM, Ho Y-J, Kii E, Alt FW. Genome-wide detection of DNA double-stranded breaks induced by engineered nucleases. Nat Biotechnol. 2015 Feb;33(2):179–186.

20. Iacovoni JS, Caron P, Lassadi I, Nicolas E, Massip L, Trouche D, et al. High-resolution profiling of gammaH2AX around DNA double strand breaks in the mammalian genome. EMBO J. 2010 Apr;29(8):1446–1457.

21. Kim D, Bae S, Park J, Kim E, Kim S, Yu HR, et al. Digenome-seq: genome-wide profiling of CRISPR-Cas9 off-target effects in human cells. Nat Methods. 2015 Mar;12(3):237–43-1 p following 243.

22. Lensing SV, Marsico G, Hänsel-Hertsch R, Lam EY, Tannahill D, Balasubramanian S. DSBCapture: in situ capture and sequencing of DNA breaks. Nat Methods. 2016 Aug;13(10):855–857.

23. Slaymaker IM, Gao L, Zetsche B, Scott DA, Yan WX, Zhang F. Rationally engineered Cas9 nucleases with improved specificity. Science. 2016 Jan;351(6268):84–88.

24. Wei P-C, Chang AN, Kao J, Du Z, Meyers RM, Alt FW, et al. Long Neural Genes Harbor Recurrent DNA Break Clusters in Neural Stem/Progenitor Cells. Cell. 2016 Feb;164(4):644–655.

25. Yan WX, Mirzazadeh R, Garnerone S, Scott D, Schneider MW, Kallas T, et al. BLISS is a versatile and quantitative method for genome-wide profiling of DNA double-strand breaks. Nat Commun. 2017;8:15058.

26. De S, Michor F. DNA secondary structures and epigenetic determinants of cancer genome evolution. Nat Struct 38 Mol Biol. 2011 Jul;18(8):950–955.

27. Moore BL, Aitken S, Semple CA. Integrative modeling reveals the principles of multi-scale chromatin boundary formation in human nuclear organization. Genome Biol. 2015 May;16(1):1270.

28. Polak P, Karlic R, Koren A, Thurman R, Sandstrom R, Lawrence MS, et al. Cell-of-origin chromatin organization shapes the mutational landscape of cancer. Nature. 2015 Feb;518(7539):360–364.

29. Whalen S, Truty RM, Pollard KS. Enhancer-promoter interactions are encoded by complex genomic signatures on looping chromatin. Nat Genet. 2016 Apr;48(5):488–496.

30. Consortium TEP. An integrated encyclopedia of DNA elements in the human genome. Nature. 2012 Sep;489(7414):57–74.

31. Chambers VS, Marsico G, Boutell JM, Di Antonio M, Smith GP, Balasubramanian S. High-throughput sequencing of DNA G-quadruplex structures in the human genome. Nat Biotechnol. 2015 Jul;33(8):877–881.

32. Rao SSP, Huntley MH, Durand NC, Stamenova EK, Bochkov ID, Robinson JT, et al. A 3D Map of the Human Genome at Kilobase Resolution Reveals Principles of Chromatin Looping. Cell. 2014 Dec;159(7):1665–1680.

33. Drier Y, Lawrence MS, Carter SL, Stewart C, Gabriel SB, Lander ES, et al. Somatic rearrangements across cancer reveal classes of samples with distinct patterns of DNA breakage and rearrangement-induced hypermutability. Genome Res. 2013 Feb;23(2):228–235.

34. Mourad R, Ginalski K, Legube G, Cuvier O. Predicting double-strand DNA breaks using epigenome marks or DNA at kilobase resolution. Genome Biol. 2018 Dec;19(1):34.

35. Canela A, Maman Y, Jung S, Wong N, Callen E, Day A, et al. Genome Organization Drives Chromosome Fragility. Cell. 2017 Jul;170(3):507–521.e18.

36. Kaiser VB, Semple CA. When TADs go bad: chromatin structure and nuclear organisation in human disease. F1000Research. 2017;6.

37. Altemose N, Miga KH, Maggioni M, Willard HF. Genomic Characterization of Large Heterochromatic Gaps in the Human Genome Assembly. PLoS Comput Biol. 2014 May;10(5):e1003628.

38. Glover TW, Wilson TE, Arlt MF. Fragile sites in cancer: more than meets the eye. Nat Rev Cancer. 2017 Aug;17(8):489–501.

39. Quint KD, Genders RE, de Koning MN, Borgogna C, Gariglio M, Bavinck JNB, et al. Human Beta-papillomavirus infection and keratinocyte carcinomas. J Pathol. 2015 Jan;235(2):342–354.

40. Zhang J, Baran J, Cros A, Guberman JM, Haider S, Hsu J, et al. International Cancer Genome Consortium Data Portal-a one-stop shop for cancer genomics data. Database. 2011 Sep;2011(0):bar026–bar026.

41. Weckselblatt B, Rudd MK. Human Structural Variation: Mechanisms of Chromosome Rearrangements. Trends Genet. 2015 Oct;31(10):587–599.

42. Viguera E, Canceill D, Ehrlich SD. Replication slippage involves DNA polymerase pausing and dissociation. EMBO J. 2001 May;20(10):2587–2595.

43. Fungtammasan A, Walsh E, Chiaromonte F, Eckert KA, Makova KD. A genome-wide analysis of common fragile sites: what features determine chromosomal instability in the human genome? Genome Res. 2012 Jun;22(6):993–1005.

44. Irony-Tur Sinai M, Kerem B. DNA replication stress drives fragile site instability. Mutat Res. 2018 Mar;808:56–61.

45. Lawrence MS, Stojanov P, Mermel CH, Robinson JT, Garraway LA, Golub TR, et al. Discovery and saturation analysis of cancer genes across 21 tumour types. Nature. 2014 Jan;505(7484):495–501.

46. Forbes SA, Beare D, Boutselakis H, Bamford S, Bindal N, Tate J, et al. COSMIC: somatic cancer genetics at high-resolution. Nucleic Acids Res. 2017 Jan;45(D1):D777–D783.

47. Wei Y, Zhang S, Shang S, Zhang B, Li S, Wang X, et al. SEA: a super-enhancer archive. Nucleic Acids Res. 2016 Jan;44(D1):D172–D179.

48. Ernst J, Kellis M. ChromHMM: automating chromatin-state discovery and characterization. Nat Methods. 2012 Feb;9(3):215–216.

49. Mrasek K, Schoder C, Teichmann A-C, Behr K, Franze B, Wilhelm K, et al. Global screening and extended nomenclature for 230 aphidicolin-inducible fragile sites, including 61 yet unreported ones. Int J Oncol. 2010 Apr;36(4):929–940.

50. Sima J, Gilbert DM. Complex correlations: replication timing and mutational landscapes during cancer and genome evolution. Curr Opin Genet Dev. 2014 Apr;25:93–100.

51. Roerink SF, Sasaki N, Lee-Six H, Young MD, Alexandrov LB, Behjati S, et al. Intra-tumour diversification in colorectal cancer at the single-cell level. Nature. 2018 Apr;556(7702):457–462.

52. Macheret M, Halazonetis TD. Intragenic origins due to short G1 phases underlie oncogene-induced DNA replication stress. Nature. 2018 Mar;555(7694):112–116.

53. Kent WJ, Sugnet CW, Furey TS, Roskin KM, Pringle TH, Zahler AM, et al. The human genome browser at UCSC. Genome Res. 2002 Jun;12(6):996–1006.

54. Grossman RL, Heath AP, Ferretti V, Varmus HE, Lowy DR, Kibbe WA, et al. Toward a Shared Vision for Cancer Genomic Data. N Engl J Med. 2016 Sep;375(12):1109–1112.

55. Delignette-Muller ML, Software CDJ of S, 2015. fitdistrplus: An R package for fitting distributions. rdrr.io.

56. Gel B, Díez-Villanueva A, Serra E, Buschbeck M, 2015. regioneR: an R/Bioconductor package for the association analysis of genomic regions based on permutation tests. academic.oup.com.

